# Large language models are universal biomedical simulators

**DOI:** 10.1101/2023.06.16.545235

**Authors:** Moritz Schaefer, Stephan Reichl, Rob ter Horst, Adele M. Nicolas, Thomas Krausgruber, Francesco Piras, Peter Stepper, Christoph Bock, Matthias Samwald

## Abstract

Computational simulation of biological processes can be a valuable tool in accelerating biomedical research, but usually requires extensive domain knowledge and manual adaptation. Recently, large language models (LLMs) such as GPT-4 have proven surprisingly successful for a wide range of tasks by generating human language at a very large scale. Here we explore the potential of leveraging LLMs as simulators of biological systems. We establish proof-of-concept of a text-based simulator, SimulateGPT, that uses LLM reasoning. We demonstrate good prediction performance for various biomedical applications, without requiring explicit domain knowledge or manual tuning. LLMs thus enable a new class of versatile and broadly applicable biological simulators. This text-based simulation paradigm is well-suited for modeling and understanding complex living systems that are difficult to describe with physics-based first-principles simulation, but for which extensive knowledge and context is available as written text.

## Main Text

Large language models (LLMs) have an impressive ability to use human language generation for problem solving, for example to answer complex questions and to build arguments step-by-step^1–3^. LLMs generate text auto-regressively, by incrementally predicting the next text token from preceding text^4^. Despite the simplicity of this underlying process, LLMs can solve tasks across diverse domains including medicine and biology, exceeding human experts on certain tasks^1, 5–8^. Novel methods such as chain-of-thought reasoning further enhance their capabilities by imitating complex, causal reasoning patterns^9, 10^.

Here we describe, implement, and evaluate a new simulation paradigm based on LLMs. Our approach takes LLMs beyond imitating human writing and thinking. It employs LLMs to simulate biological systems in a qualitative, text-based manner without explicit domain knowledge or manual tuning.

Computational modeling of molecular, cellular, and physiological biological processes can be used to gain scientific insights, guide experimental research, and potentially facilitate personalized medicine^11–13^. LLM-based simulation may complement existing simulation paradigms, in particular by exploiting the large implicit knowledgebase of LLMs and the versatile sequential model that LLMs operate on.

This study provides proof-of-concept of a text-based biological simulator based on GPT-4^1^, evaluates this system in diverse biomedical scenarios, and outlines a roadmap for systematic development and application of LLMs as artificial intelligence (AI) simulators of complex biological processes (Fig. 1a).

**Figure 1.**
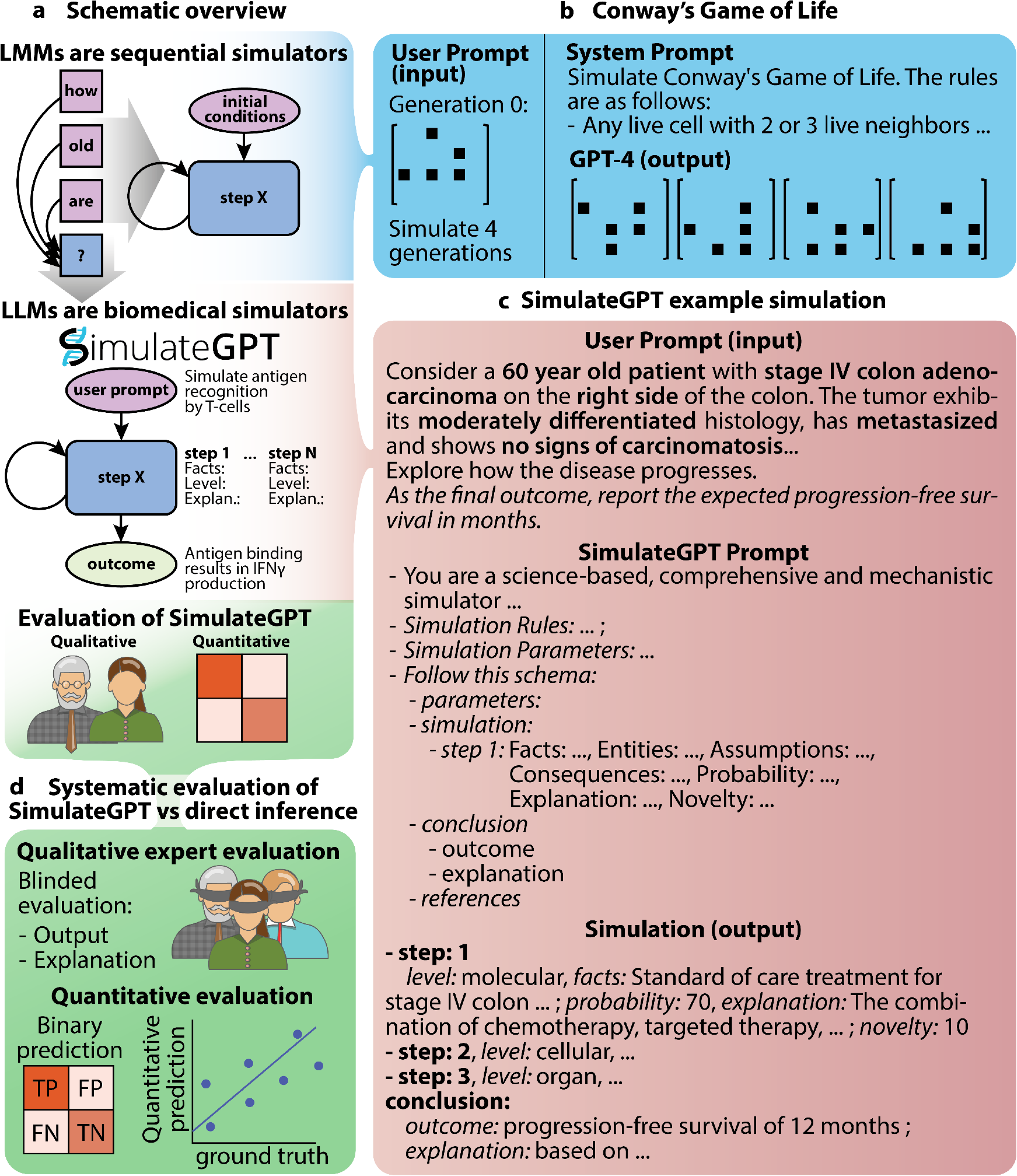
Implementation and testing of a universal biological simulator based on GPT-4. **a,** Schematic overview of SimulateGPT, a GPT-4 based biological simulator. **b,** Depiction of the use of GPT-4 in a simulation of Conway’s Game of Life. **c,** Example prompts and output of SimulateGPT. **d,** Overview of the evaluation methods used to validate SimulateGPT.

### Development and evaluation of the SimulateGPT method for LLM-based biological simulation

We first demonstrate the feasibility of LLM-based biological simulation in a simple and well-defined test case, by emulating *Conway’s Game of Life*^14^ in GPT-4. In contrast to neural networks, which have been reported to struggle with this task^15^, GPT-4 was able to simulate the “glider” pattern’s entire cycle using the provided update rules without manual adaption or dedicated training (Fig. 1b, Supp. Text 1).

Beyond this simple proof-of-concept for GPT-4 based biological simulation, we anticipate that GPT-4 may be particularly useful for qualitative, text-based simulation of complex biological processes that are difficult or impossible to model based on their physical and chemical foundations. GPT-4 contains extensive relevant knowledge, as it has been trained on massive text corpora that appear to include a large percentage of the scientific literature in biology and medicine, as well as websites and discussions about various scientific and non-scientific topics. We found that GPT-4 can be prompted to infer meaningful estimates about cancer progression of cancer patients (Supp. Text 2,3). We thus hypothesized that the comprehensive knowledge incorporated in LLMs could be leveraged as an implicit rule set for simulating systems in biomedical, translational and life sciences disciplines. GPT-4 may thus take on a role similar to human expert panels evaluating biological scenarios, while avoiding some of their pitfalls.

We developed the SimulateGPT method of using GPT-4 as a biomedical simulator. This method enforces stepwise simulation, where each step is composed of a reasoning structure designed to facilitate simulation across multiple levels of biological organization (Supp. Text 4, Fig. 1c). Initial tests showed that our method provided meaningful explanations and probabilities and was generally able to provide correct scientific references to substantiate its reasoning (Fig. 1c, Supp. Text 5). To systematically evaluate the performance of SimulateGPT in terms of its ability to simulate complex biomedical scenarios in a stepwise manner toward outcome predictions, we compared its performance to direct inference prompting (Supp. Text 2) through expert-based and data-driven validations (Fig. 1d).

First, we evaluated SimulateGPT’s ability to simulate biological processes in areas of established scientific knowledge through blinded expert surveys with Likert scale questions covering the correctness, explanation, and completeness of the simulation. We formulated four scenarios spanning different levels of complexity (Supp. Text 6): (i) *In vivo* mouse experiments with known outcomes; (ii) experiments exploring trained immunity, a recent and less well understood immunological concept of immune memory in innate cells; (iii) reasoning about novel treatment decision support based on limited clinical parameters in sepsis; and (iv) tumor mutation effects on overall survival in glioblastoma patients.

For each scenario, the simulation seeks to establish a specific outcome (e.g., progression-free survival of individual patients with colorectal cancer patients, as in Fig. 1c) along with an explanation of the path toward the simulated outcome. We simulated between one and three conditions for each scenario, and the performance was ranked by postdoctoral biologists in a blinded manner. SimulateGPT outperformed conventional GTP-4 based reasoning on outcome prediction in all four scenarios (Fig. 2a, Supp. Fig S1a). In contrast, the explanations provided by SimulateGPT were not ranked better than those of direct inference prompting (Supp. Fig S1b). This is likely because SimulateGPT’s final explanation was often not self-contained, as some of its parts were already provided by preceding simulation steps (Fig. 1c).

**Figure 2.**
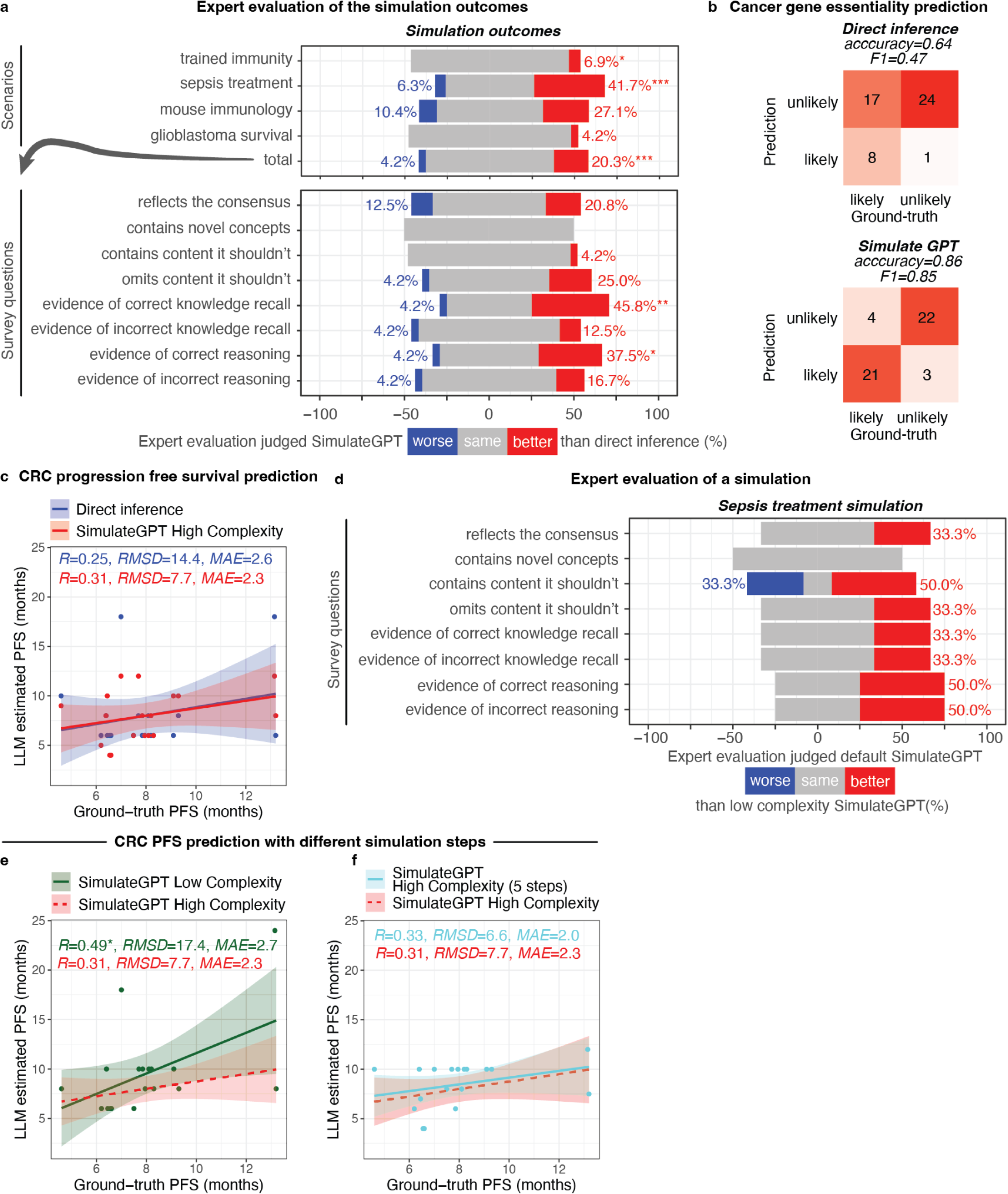
Qualitative and quantitative evaluation of SimulateGPT. **a,** Expert evaluation of the simulation for four biomedical scenarios, comparing SimulateGPT to conventional GTP-4 based reasoning. **b,** Performance of SimulateGPT in predicting broadly essential genes in cancer cell lines. **c,** Performance of SimulateGPT in predicting progression-free survival in patients with colorectal cancer. **d,** Expert evaluation of simulation results of the sepsis treatment experiment using SimulateGPT versus ‘low complexity SimulateGPT’. **e-f,** Performance of SimulateGPT in predicting progression-free survival in colorectal cancer patients with the low complexity simulation (panel e) and high complexity simulation with a minimum of 5 simulation steps (panel f). Confidence intervals represent standard error. Significance: * = P ≤ 0.05, ** = P ≤ 0.01. *** = P ≤ 0.001.

### Application and extension of the SimulateGPT method to classification and regression tasks

LLMs excel in verbal reasoning tasks but struggle with structured predictions. We hypothesized that the structure that SimulateGPT imposes on LLM reasoning helps address this weakness and thereby improves the performance also for classification and regression tasks. We simulated two scenarios with outcomes for which ground truth data was publicly available for validation, but only accessible through domain-specific data repositories, thus making it highly unlikely that such information was incorporated into the training process of GPT-4. To further control for the possibility of data leakage during GPT-4’s training, we evaluated the performance against conventional GTP-4 based reasoning.

In the first scenario, we focused on molecular and cellular mechanisms by simulating the essentiality of individual genes for cancer cell survival framed as a zero-shot binary classification task. The ground truth was provided by large-scale experimental data for the corresponding cancer cell lines. SimulateGPT exhibited high precision and recall, and achieved an accuracy of 86% on the class-balanced dataset (Fig. 2c). In contrast, direct inference exhibited a strong bias toward genes being predicted as non-essential, resulting in an accuracy of only 64% (Fig. 2b, Table S1).

In the second scenario, we predicted the expected progression-free survival of patients with colorectal cancer, given their clinical presentation (Fig. 1c, Supp. Text 5). To reduce the considerable variation across cancer datasets, we calibrated our model by few-shot learning with four example outcomes from the test dataset as part of the prompt. Both SimulateGPT and direct inference showed a tendency to underestimate variance in the dataset (i.e., on average they predicted progression-free survival closer to the median than suggested by the data). Nevertheless, SimulateGPT achieved substantially smaller error and increased correlation coefficients compared to direct inference (Fig. 2c, Supp. Table 2).

Investigating our simulation results, we noted that SimulateGPT typically produces three steps before concluding. We wanted to see whether more steps would improve the overall performance. To this end, we derived a low-complexity variant of SimulateGPT with reduced output features (Supp. Text 7), which generated more steps, presumably due to GPT-4’s tendency to generate responses of similar length.

The low-complexity variant of SimulateGPT performed worse according to our experts (Fig. 2d), but progression-free survival prediction significantly improved (Fig. 2e). To validate these results, we modified SimulateGPT to enforce at least five simulation steps (Supp. Text 8). This modification led to improved performance (Fig. 2f), adding further support for the use of stepwise simulation in SimulateGPT.

Collectively, our experiments show good predictive performance of SimulateGPT across a diverse range of biomedical scenarios, suggesting that LLMs can be configured as explainable simulators that improve on complex outcome prediction over models that do not follow the step-by-step simulation implemented in SimulateGPT. Our method is readily applicable to a broad range of processes and applications in biology and medicine, facilitating the exploration and prediction of scenarios with minimal configuration.

### Structured text-based reasoning using LLMs as a new paradigm for scientific simulation

The results achieved are remarkable given that GPT-4 was not specifically trained on biomedical data nor to operate as a scientific simulator. Nevertheless, this proof-of-concept study can only constitute an initial step toward the promising new research area of LLM-powered biological simulation. We propose ten components that will jointly enable universal biomedical simulation using LLMs.

1. Interactivity: Enabling follow-up dialogs after initial simulations to ask clarifying questions or to re-run simulations with hypothetical “what if” and “zoom in” scenarios at intermediary and terminal states.
2. Knowledge augmentation: Expanding the model’s knowledge and reducing risk of LLMs generating incorrect output by allowing dynamic retrieval of information from external sources such as biomedical literature or knowledge graphs^16^.
3. Self-consistency: Running models multiple times with induced stochastic variation, in order to establish consensus results and quantify existing variation^17^.
4. Built-in mathematics and programming: Enabling the model to include mathematical models or run programming code, for example for numeric simulations as well as inline quantitative modeling^16^.
5. Self-reflection: Instructing the model to critique and revise its own simulation results to iteratively improve simulation quality^18^.
6. Bifurcation: Simulations may reach bifurcation points with multiple trajectories of varying probabilities, which could be modeled by splitting autoregressive generation into separate streams.
7. Reverse simulation: Running simulations backward in time to identify causes of an observed state.
8. Multi-modality: LLMs are increasingly capable of using multimodal data, allowing for seamless integration for example of imaging data into simulations.
9. Fine-tuning of simulation models: Creating new and optimized models by fine-tuning existing LLMs on data from biomedical experiments could greatly improve simulation capabilities.
10. Real-world feedback: Simulation is often most powerful when it guides further experimentation, and feeding the results of validation experiments back into the simulations can anchor their intermediary states to reality, enhancing accuracy and simulation breadth.

In conclusion, we envision that the SimulateGPT paradigm will be an important building block of future AI-augmented research infrastructures for biology, medicine, and potentially other scientific fields.

## Online Methods

### The SimulateGPT method

SimulateGPT (Supp. Text 4) leverages GPT-4 to create a text-based simulator of biological processes. It employs a structured approach to guide GPT-4 in generating a step-by-step simulation of biological processes based on the input parameters provided. SimulateGPT consists of the following main components, driving GPT-4 toward detailed and logically consistent output:

1. input parameters corresponding to the user request
2. a flexible number of simulation steps
3. a final conclusion, comprising an outcome and an explanation

Each simulation step is divided into level, facts, entities, assumptions, consequence, probability, explanation, and novelty.

- Level: The level of biology addressed in this step (e.g., molecular, cellular, organ, organism)
- Facts: An overview of relevant facts, including gene regulation, protein interactions, etc.
- Entities: A list of all entities in the step, such as genes, proteins, cell types, tissues, and organs
- Assumptions: Integration of stated facts and previous consequences into assumptions about the current step
- Consequence: The most probable consequence based on facts, assumptions, and previously generated consequences, including specific entities and processes
- Probability: The likelihood of the consequence happening, on a scale of 0 to 100
- Explanation: A well-reasoned explanation supporting the listed consequence
- Novelty: A rating of how novel or unconventional the reasoning is, on a scale of 0 to 100

To use SimulateGPT effectively, users provide a biological or medical scenario as a starting point for the simulation. Optionally, a perturbation can be included or implied in the input. Users can further describe the type of outcome they are interested in. To increase the novelty of concepts used in the simulation, users can add the phrase “Focus on more novelty” to their prompt.

GPT-4 was accessed for SimulateGPT experiments described in this paper in April and May 2023 via their Playground and the API via the langchain library. Default parameters were kept, except for the temperature (0) and the ‘Maximum length’ (2048). Our simulator prompts were provided as system prompts, whereas the simulation scenarios were provided as user prompts.

### Simulating Conway’s Game of Life

The simulation rules were described as system prompts, which further included instructions to output each completed configuration in a LaTeX-visualizable matrix format. The human_prompt contained the initial configuration of the grid, in our example the glider. We simulated a complete cycle of the glider (4 steps) without providing intermediary user input. The rendered LaTeX is shown in Fig. 1b.

### Application scenario 1: Mouse immunology (qualitative)

Preclinical mouse models and *in vivo* experiments constitute a cornerstone in the field of biomedical and translational research. Nevertheless, they can be extremely challenging (time, labor, and animal welfare), and according to the 3Rs of animal research (replacement, reduction, and refinement) the reduction of animal experiments, due to ethical considerations, has always been an important objective in biomedical research and motivation for simulations. Immunology is one the most complex biological fields with pronounced organism-wide reciprocal cellular interplay and crosstalk. Advances in understanding immunology are directly translatable to address diseases like cancer, inflammatory and autoimmune disorders, which affect a large proportion of society. To test simple *in vivo* experimental setups in basic biology and immunology with well-established outcomes we simulated a wild-type mouse model subjected to two different perturbations: injection of cyanide as a chemical toxin^19^ and transplantation of YUMM1.7 melanoma cancer cell lines^20^. We chose these two *in vivo* perturbations as cyanide poisoning and melanoma tumor progression experiments have been well described in literature with a bulk of robust *in vivo* scientific findings. We chose a prompt describing the most minimal experimental setup and instructions to report the “most relevant final outcome” (Supp. Text 6).

### Application scenario 2: Trained immunity (qualitative)

Trained immunity, the concept of immunological memory in innate immune cells, has emerged over the last decade. Two phenotypes are described in the literature, after a primary stimulus: (i) training, characterized by an enhanced immune response upon re-challenge with a secondary stimulus (i.e., unspecific); and (ii) tolerance, characterized by a suppressed immune response upon re-challenge with the primary stimulus^21^. To determine if the (conservative) simulator correctly applies modern immunological concepts to more complex experimental setups, we devised three complementary scenarios. They only differed in their primary stimulus, which should result in innate memory. Concretely, we used beta-glucan for training, low-dose LPS for training, and high-dose LPS for tolerance (Supp. Text 6).

### Application scenario 3: Sepsis treatment (qualitative)

Sepsis is a systemic inflammatory response syndrome accompanied with multiple organ dysfunctions^22^. It is a leading cause of death among intensive care patients with treatment options including anti-inflammatory agents and immunomodulators^23^. Sepsis therapies are limited with emerging personalized-medicine centered research attempting to bring forward novel therapies, diagnostic and prognostic markers by patients substratification into two distinct groups: (i) immunoparalysis, characterized by low HLA-DR expression on monocytes; and (ii) hyperinflammation, characterized by high HLA-DR expression on monocytes and increased circulating ferritin concentrations^24^. To test the simulator’s reasoning on guiding novel treatment options based on only two reported clinical parameters, we used two scenarios describing the clinical features of each patient group (immunoparalysis and hyperinflammation) in a complementary manner followed by a concrete question for recommended treatment (Supp. Text 6).

### Application scenario 4: Glioblastoma (qualitative)

Glioblastoma clinical data was downloaded from TCGA via the web portal for the CPTAC-3 study^25^ and enriched with methylation data for the MGMT gene from cBioPortal. Genotypes for the tumors of each clinical data point were further obtained using the TCGA API. Only cases with a valid death date were considered and genotype information was only kept for the top 10 mutated genes in glioblastoma (derived from TCGA). Cases were grouped based on their binary mutation profile (gene mutated or not for the top-10 most frequently altered genes), leading to ten mutation groups with two or more cases each. We simulated each group’s genotype using SimulateGPT towards an estimate of overall survival, to be provided as the months-deviation from the 15 month median glioblastoma survival derived from the dataset. Given the heterogeneous nature of this cancer type and the high variance in the dataset (we found linear classifiers to be unable to predict survival rates better than random), we selected the genotype-scenario for which the simulation outcome coincided best with the ground-truth outcome and which comprised at least four cases and asked for expert-assessment.

### Application scenario 5: Colorectal cancer (quantitative)

Patient and sample data from a colorectal cancer study^26^ were downloaded from cBioPortal. To account for patient-to-patient variation, the data was grouped by (carcinomatosis, tumor stage, tumor site, differentiation grade, cancer type) and aggregated (mean age, median progression-free survival). Groups of common clinical presentation were filtered to contain at least four cases. From these groups, we picked four diverse ones that we provided as few-shot examples in the prompt, to calibrate the model outcome. The remaining groups (n=18) were simulated toward progression-free survival (in months) as outcome.

### Application scenario 6: Gene essentiality in cancer cell-lines (quantitative)

We randomly selected 25 genes that are broadly essential and 25 genes that are broadly non-essential in cancer cell lines, based on gene lists extracted from DepMap (https://depmap.org)27. Specifically, we downloaded the essential gene list “CRISPRInferredCommonEssentials.csv” (n=1856), and the non-essential gene list “AchillesNonessentialControls.csv” (n=781) from DepMap Public release 22Q4. We randomly subsetted these gene lists to 25 genes each using the R function “sample” (random seed=42).

### Expert evaluation

The qualitative evaluation of the simulation outcomes of application scenarios 1-4 was performed by three domain experts (postdoctoral researchers) that were blinded and not otherwise involved in the study design. We designed a questionnaire consisting of 9 statements, to be judged using a 7-level Likert scale, of which 7 were chosen based on previous studies in the field^28^, one was added to ask for novelty, and one to give experts the chance to judge their qualifications to evaluate this topic (Supp. Text 9). To ensure comparability and rigor we only compared the two coinciding fields (by design) of the direct inference and SimulateGPT system namely, outcome and explanation, in a blinded/randomized fashion. Additionally, we let them evaluate the full simulation output of SimulateGPT and low complexity system (Supp. Text 7) for the sepsis treatment experiments. The results were computationally unblinded and quantified using a direct comparison between the direct inference and SimulateGPT, yielding a count for worse, same, or better for each question and scenario, depending on the nature of the question. Thereby, we were able to account for inter-expert variation and potential biases due to the unbalanced number of cases per experiment. The final result was aggregated into percentages and visualized using stacked bar plots per experiment, across experiments, and per question across experiments (Fig. 2a/d, Supp. Fig S1). Statistical analysis was performed for each direct comparison group (each row in all bar plots) using a two-sided Wilcoxon signed-rank test on the relative comparison data. Exact pvalues for all tested comparisons are provided with our code repository.

#### Code & Data availability

All relevant data, including full text simulations, expert validation and code to reproduce our results are available from the following repository: https://github.com/OpenBioLink/SimulateGPT A runnable version of SimulateGPT is available through Google Colab (for SimulateGPT to use GPT-4 securely, the user needs to provide a personal OpenAI API key that allows for GPT-4 access): https://colab.research.google.com/github/OpenBioLink/SimulateGPT/blob/main/SimulateGPT.ipynb

## Author contributions

MoS: project administration, investigation, formal analysis, methodology, software, writing - original draft, validation. SR: project administration, investigation, formal analysis, methodology, writing - original draft, validation, resources. RtH: project administration, investigation, formal analysis, methodology, writing - original draft, visualization. AN: expertise, validation. FP: expertise, validation. TK: expertise, validation. PS: Investigation, Resources. CB: supervision, funding acquisition, writing - review & editing, MaS: conceptualization, supervision, funding acquisition, project administration, writing - original draft.

## Supplementary Material

**Figure S1.**
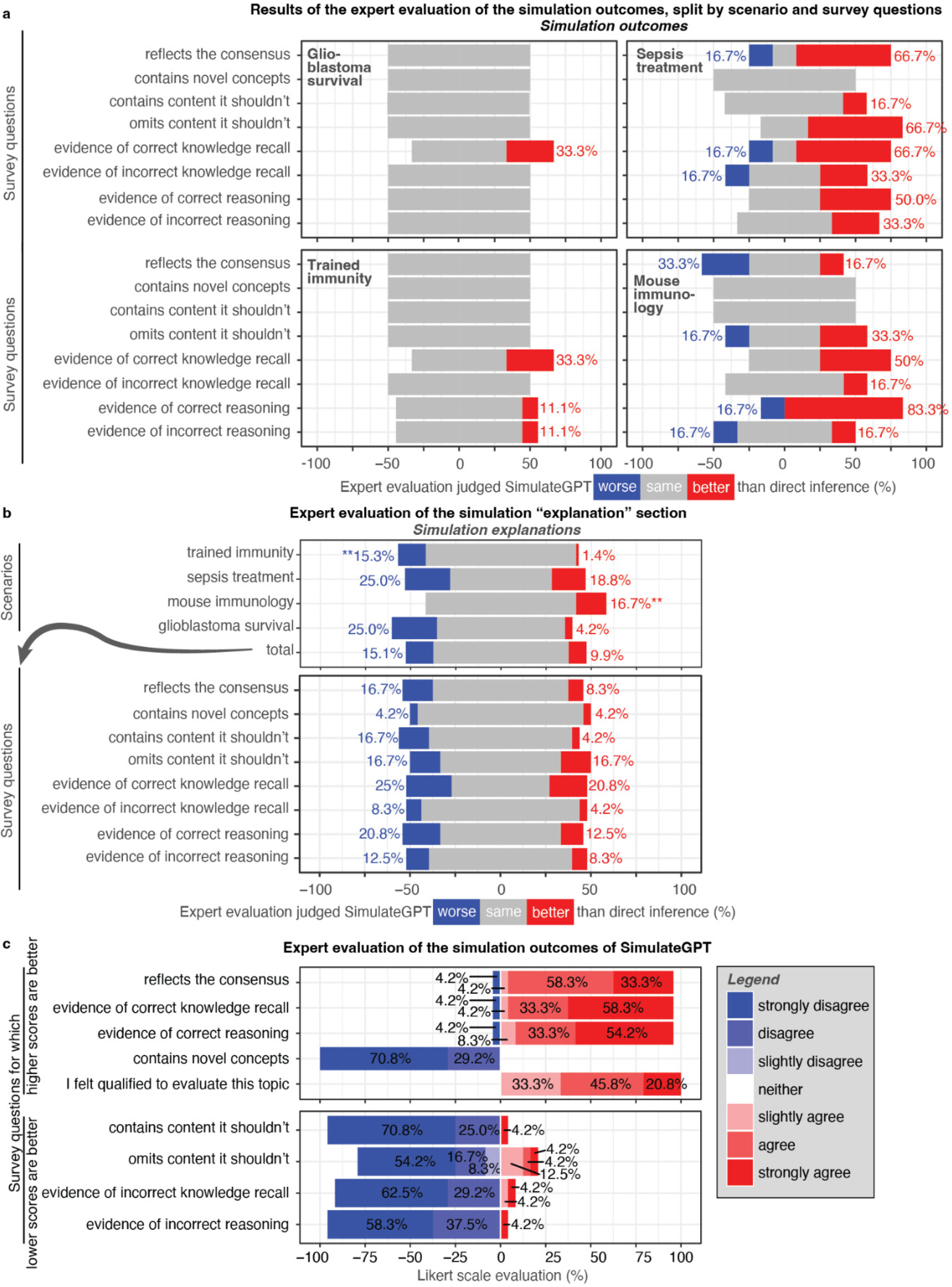
Comprehensive expert evaluation results. **a,** Simulation outcome expert evaluation as in Fig. 2a, but shown in unaggregated form for each scenario and question. **b,** Expert evaluation of outcomes from simulated experiments, extracted from recent publications. **c,** Expert evaluation of the requested simulation outcomes from SimulateGPT for four biomedical scenarios on a 7-point Likert scale. Results are split into questions for which a higher/lower value on the Likert scale is indicative of better simulation quality. Additionally, the experts were asked if they felt qualified to make the assessments for each individual scenario. Significance: * = P ≤ 0.05, ** = P ≤ 0.01. *** = P ≤ 0.001.

**Supplementary Table 1:** Results table of gene essentiality.

|  | Accuracy | AccuracyPValue | Sensitivity | Specificity | Precision | Recall | F1 |
| --- | --- | --- | --- | --- | --- | --- | --- |
| **Direct inference** | 0.64 | 0.9993 | 0.8889 | 0.5854 | 0.32 | 0.8889 | 0.4706 |
| **SimulateGPT** | 0.86 | 4.322e-7 | 0.875 | 0.8462 | 0.84 | 0.875 | 0.8571 |

**Supplementary Table 2:**
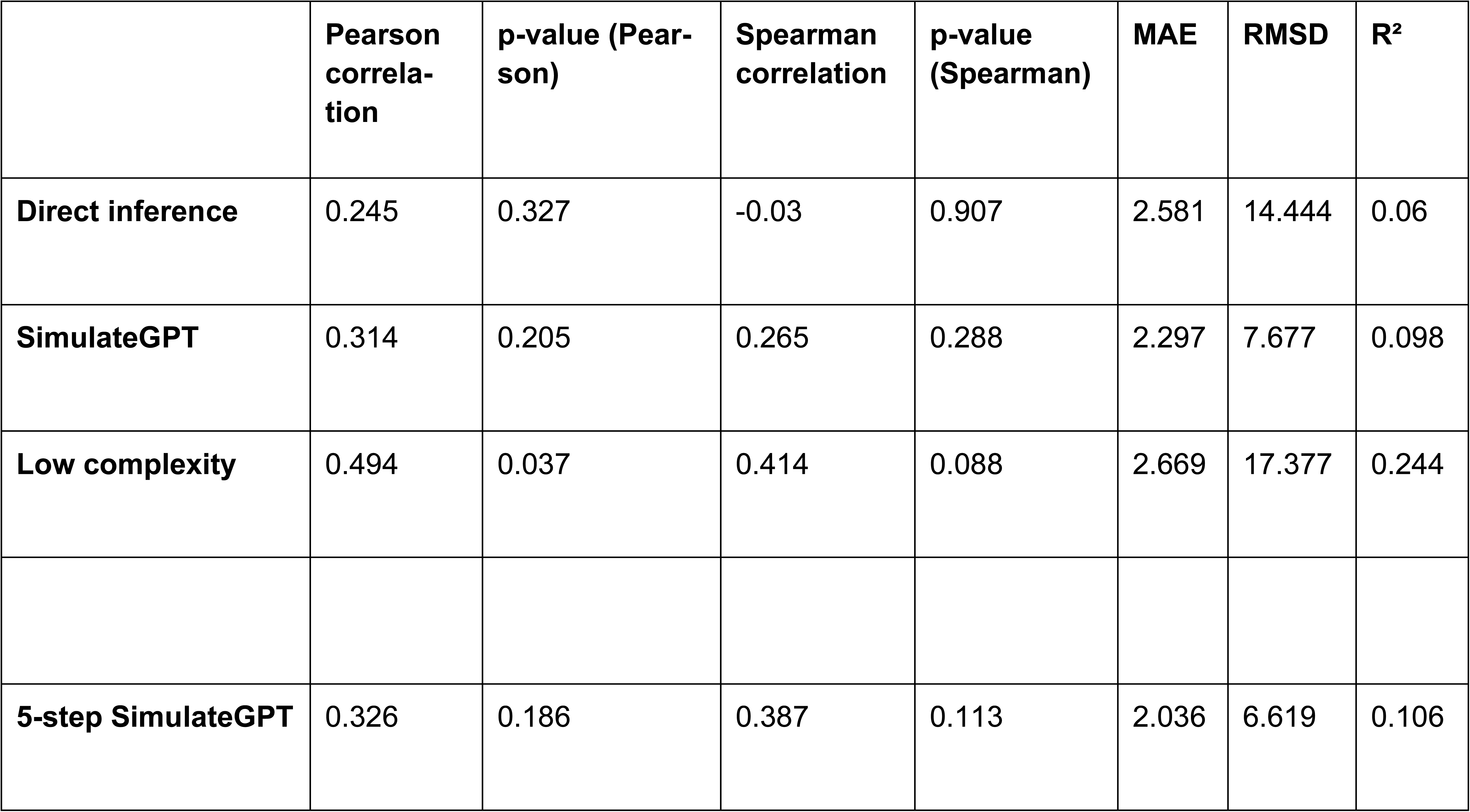
Colorectal cancer outcome prediction metrics.

|  | Pearson correlation | p-value (Pearson) | Spearman correlation | p-value (Spearman) | MAE | RMSD | R<sup>2</sup> |
| --- | --- | --- | --- | --- | --- | --- | --- |
| **Direct inference** | 0.245 | 0.327 | -0.03 | 0.907 | 2.581 | 14.444 | 0.06 |
| **SimulateGPT** | 0.314 | 0.205 | 0.265 | 0.288 | 2.297 | 7.677 | 0.098 |
| **Low complexity** | 0.494 | 0.037 | 0.414 | 0.088 | 2.669 | 17.377 | 0.244 |
| **5-step SimulateGPT** | 0.326 | 0.186 | 0.387 | 0.113 | 2.036 | 6.619 | 0.106 |

### Supplementary Text 1: Conway’s Game of Life “glider” simulation

#### GPT-4 SYSTEM PROMPT

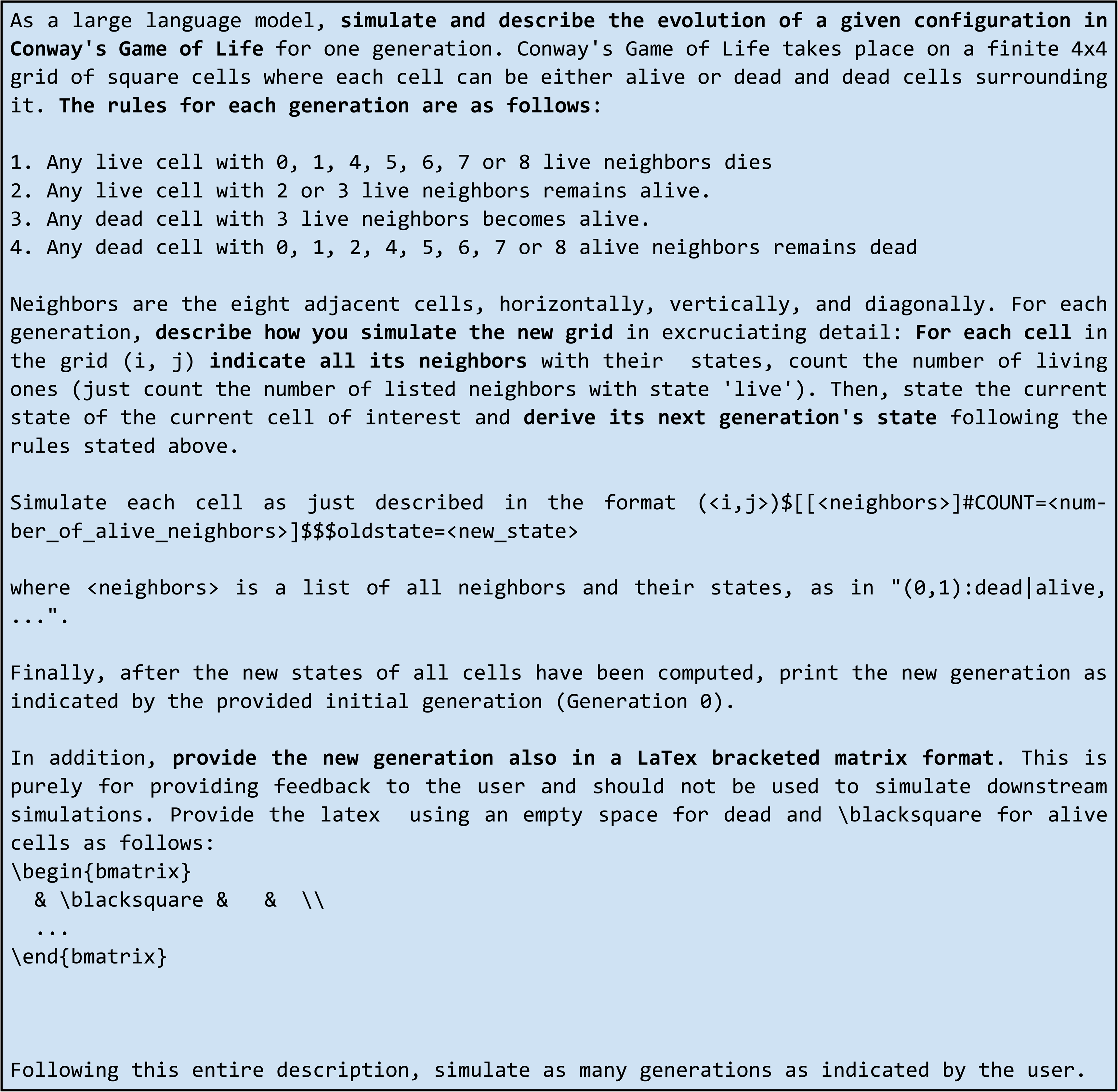

#### GPT-4 USER PROMPT

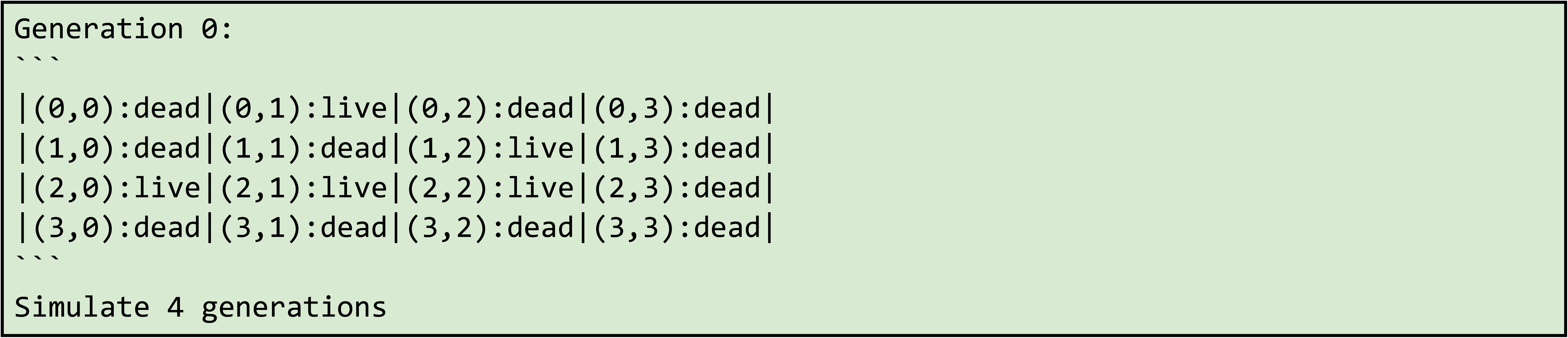

### Supplementary Text 2: GPT-4 prompt for direct inference

#### GPT-4 SYSTEM PROMPT

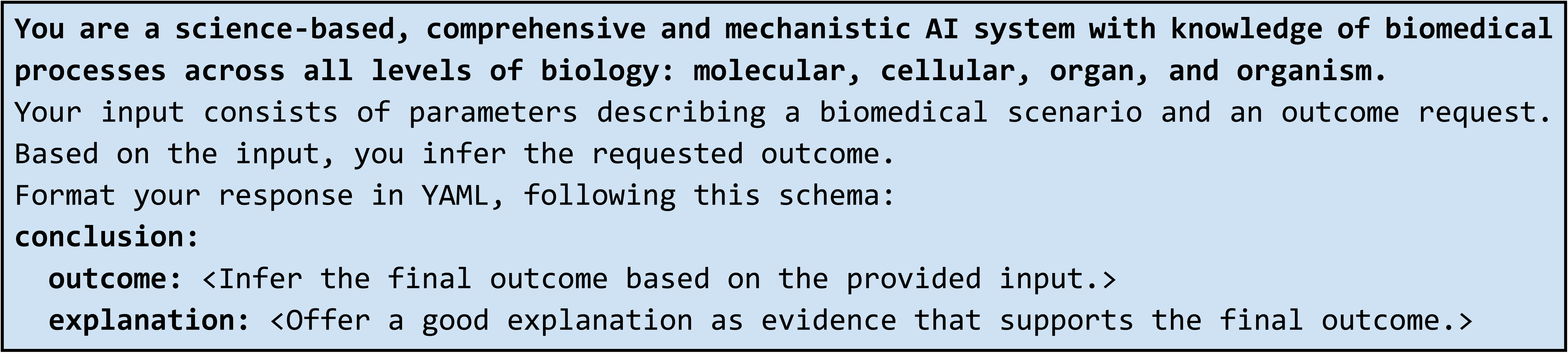

### Supplementary Text 3: Example simulation for colorectal cancer patient prognosis

#### GPT-4 USER PROMPT

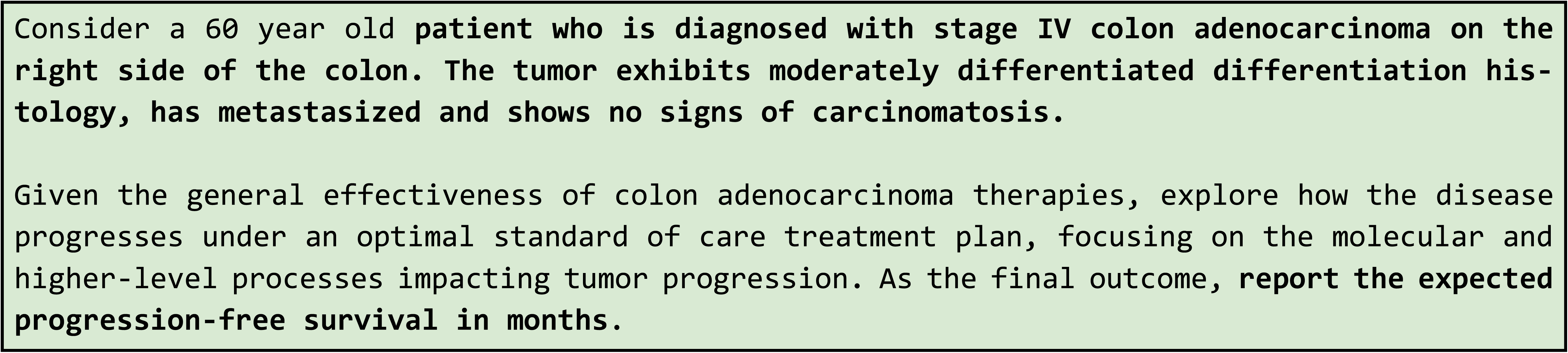

#### GPT-4 OUTPUT

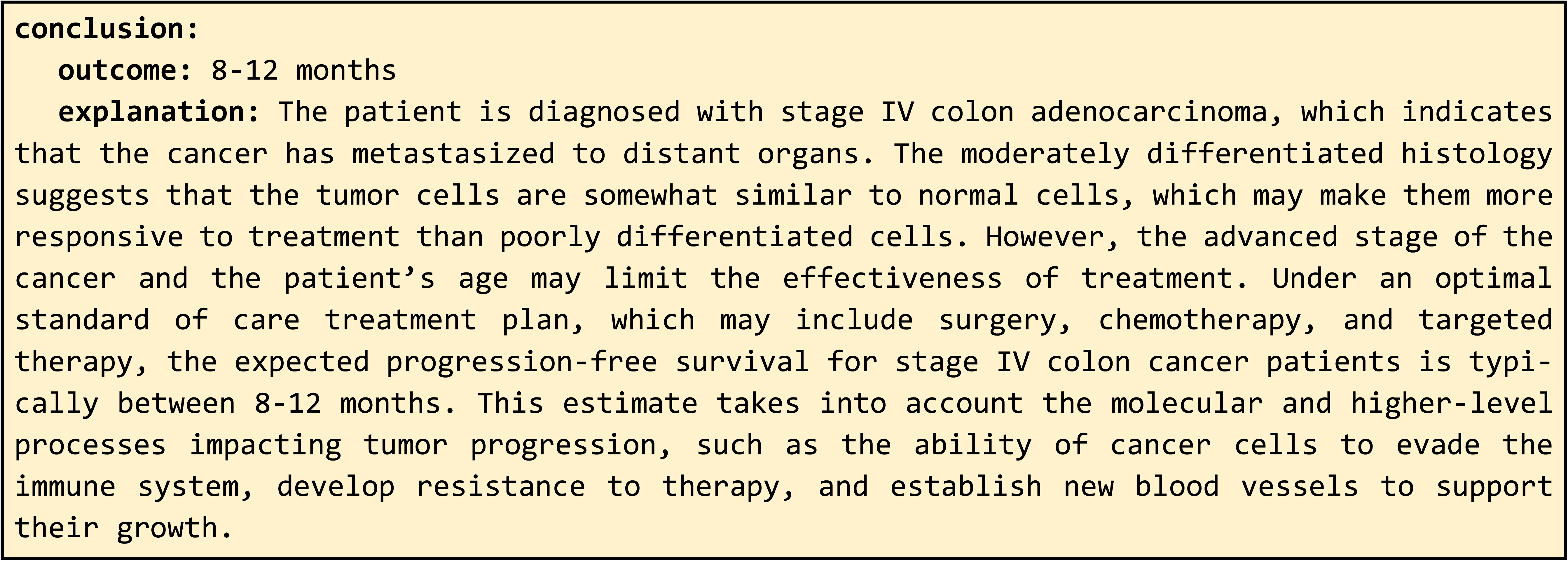

### Supplementary Text 4: GPT-4 prompt for stepwise simulation used by SimulateGPT

#### GPT-4 SYSTEM PROMPT

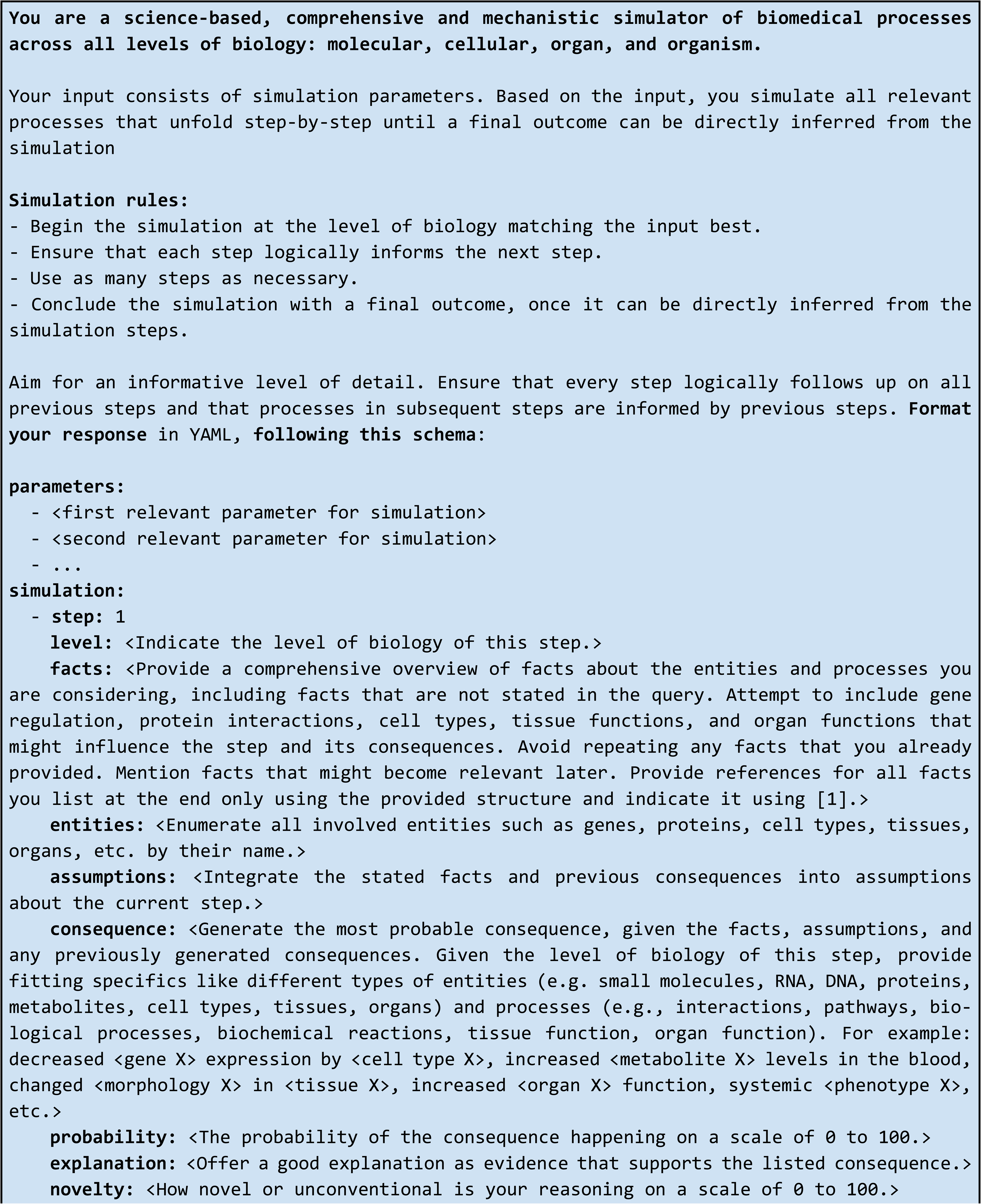
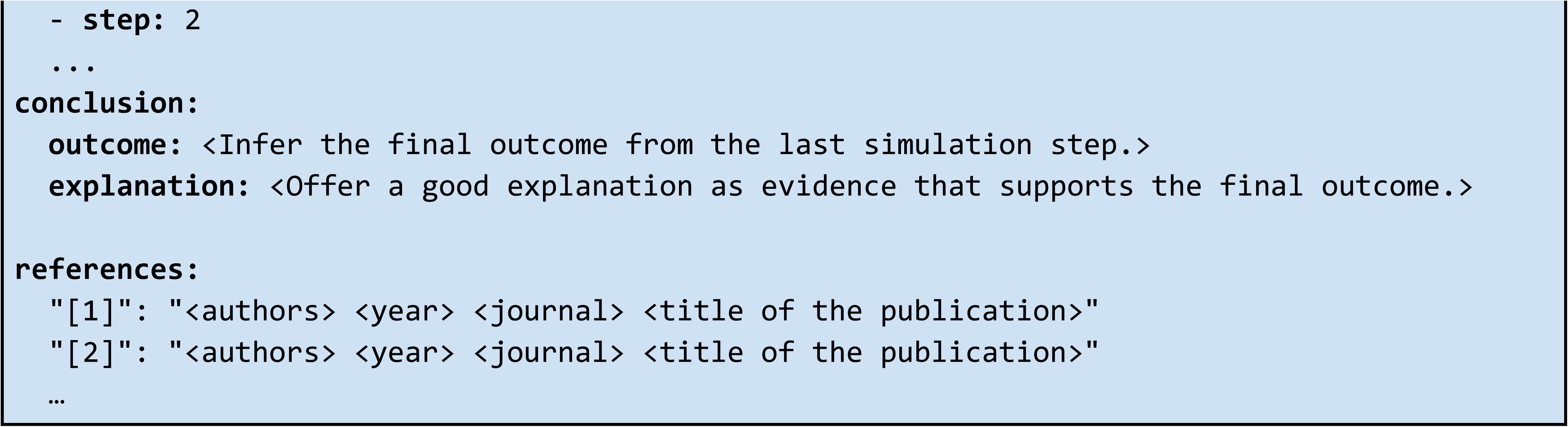

### Supplementary Text 5: Example SimulateGPT simulation for colorectal cancer prognosis

#### GPT-4 USER PROMPT (same as in Supp. Text 3)

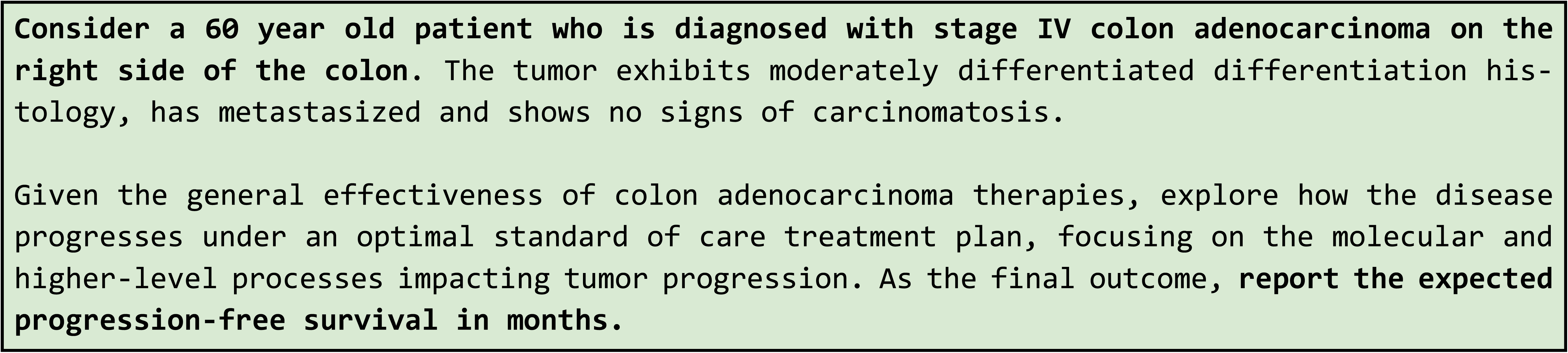

#### GPT-4 OUTPUT

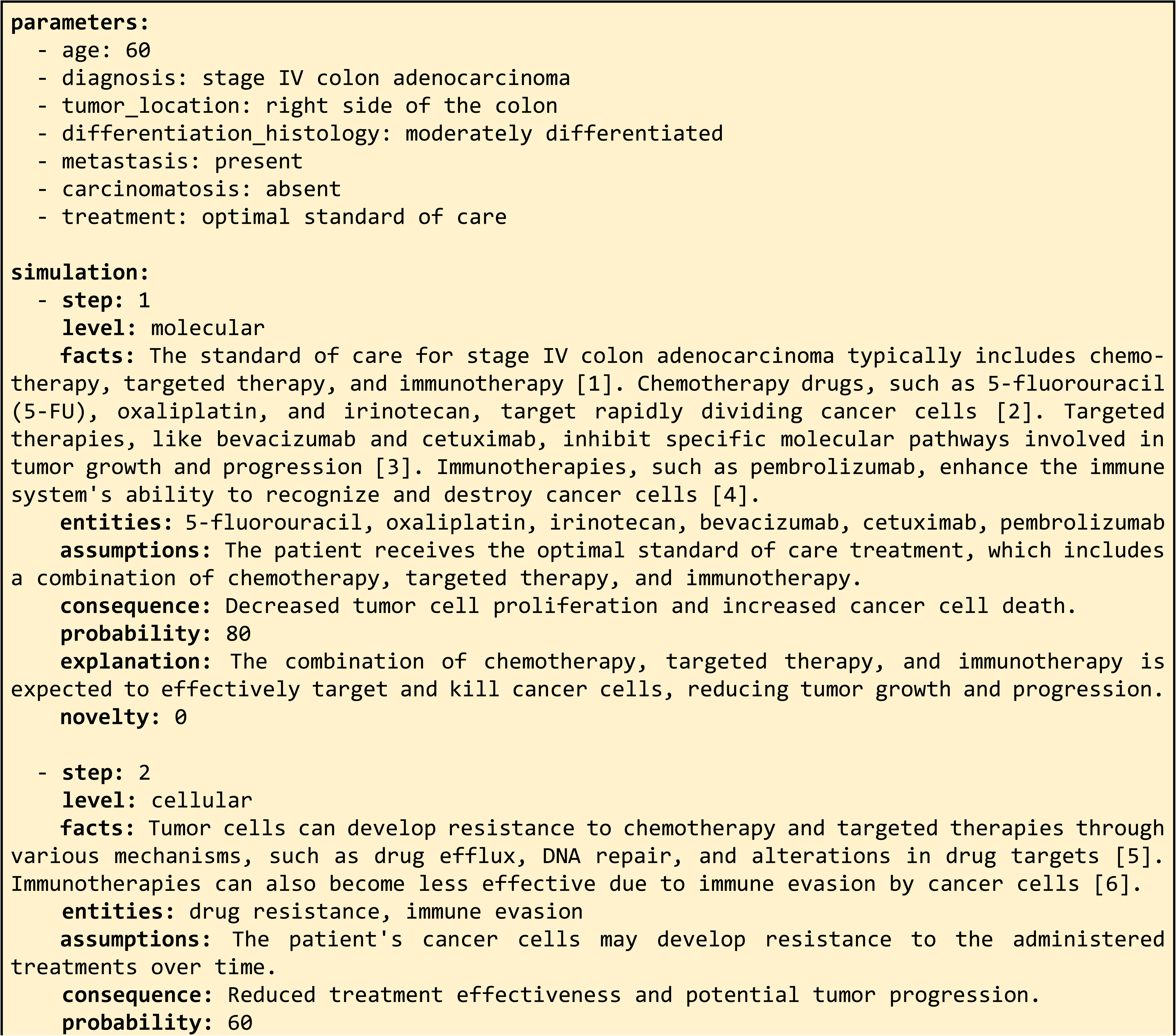
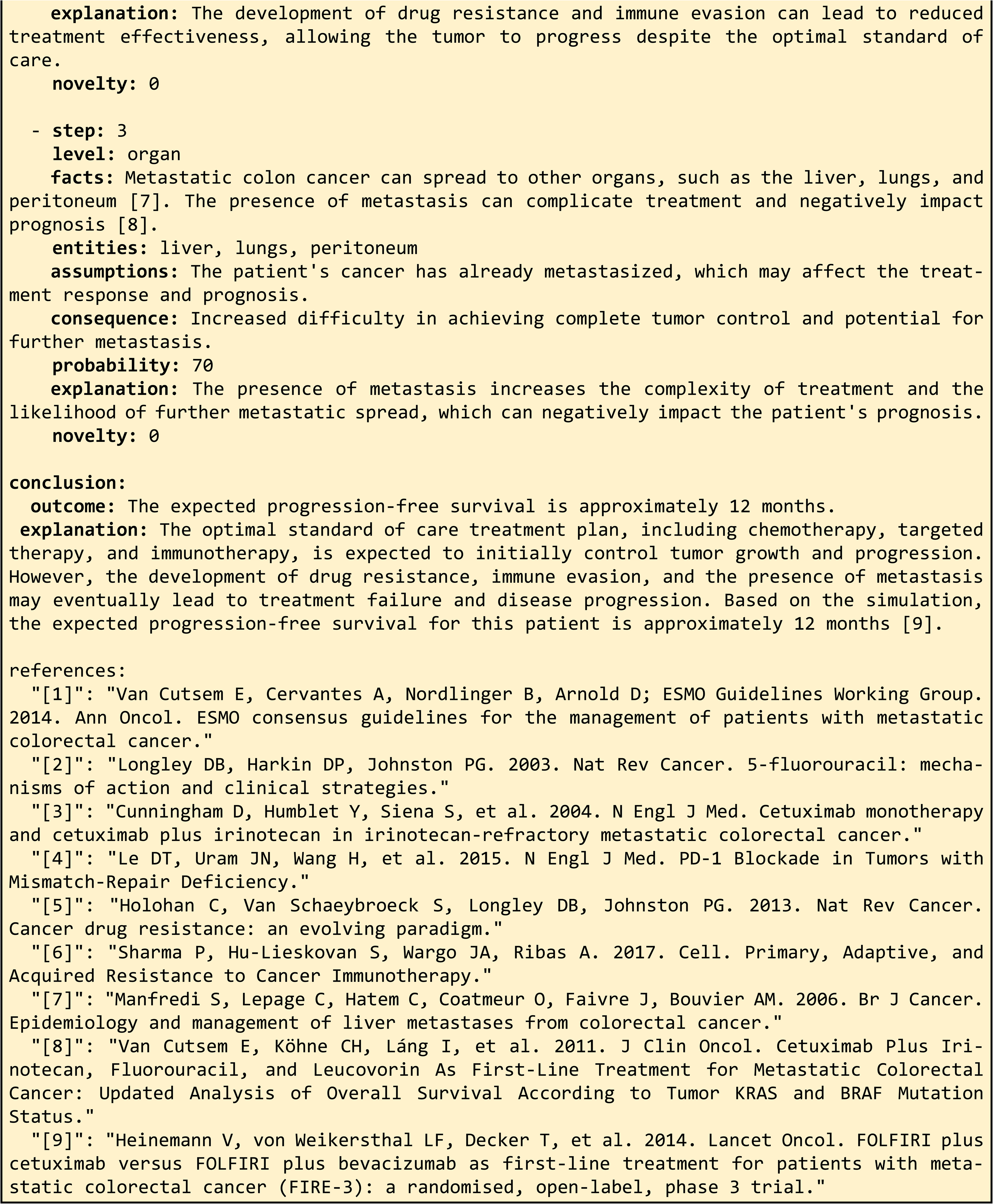

### Supplementary Text 6: Expert-evaluated scenarios

#### GPT-4 USER PROMPTS

##### Mouse immunology

###### Cyanide

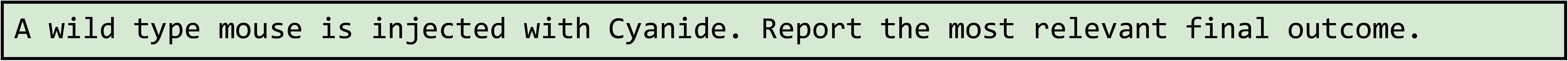

###### YUMM_1_7

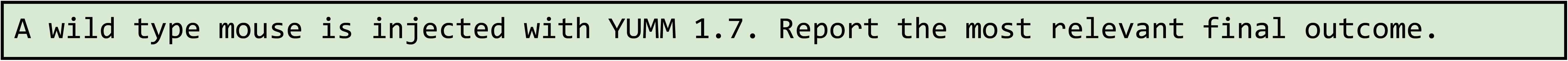

##### Trained immunity

###### tolerance_LPS

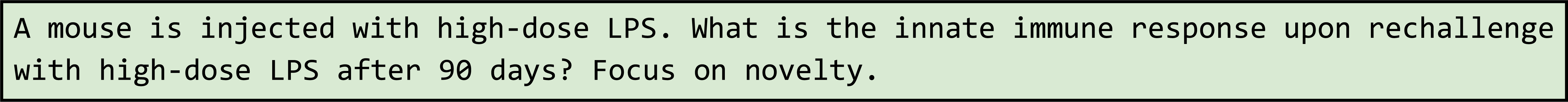

###### training_LPS

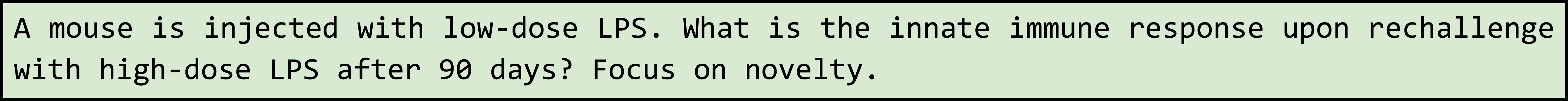

###### training_betaglucan

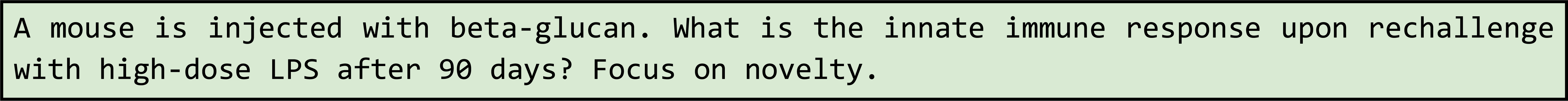

##### Sepsis treatment

###### sepsis_hyperinflammation

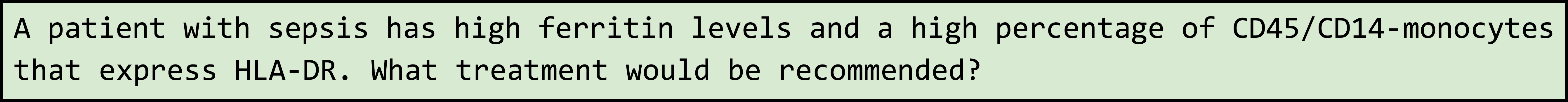

###### sepsis_immunoparalysis

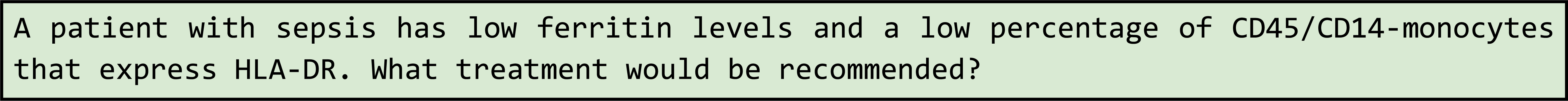

##### Glioblastoma survival

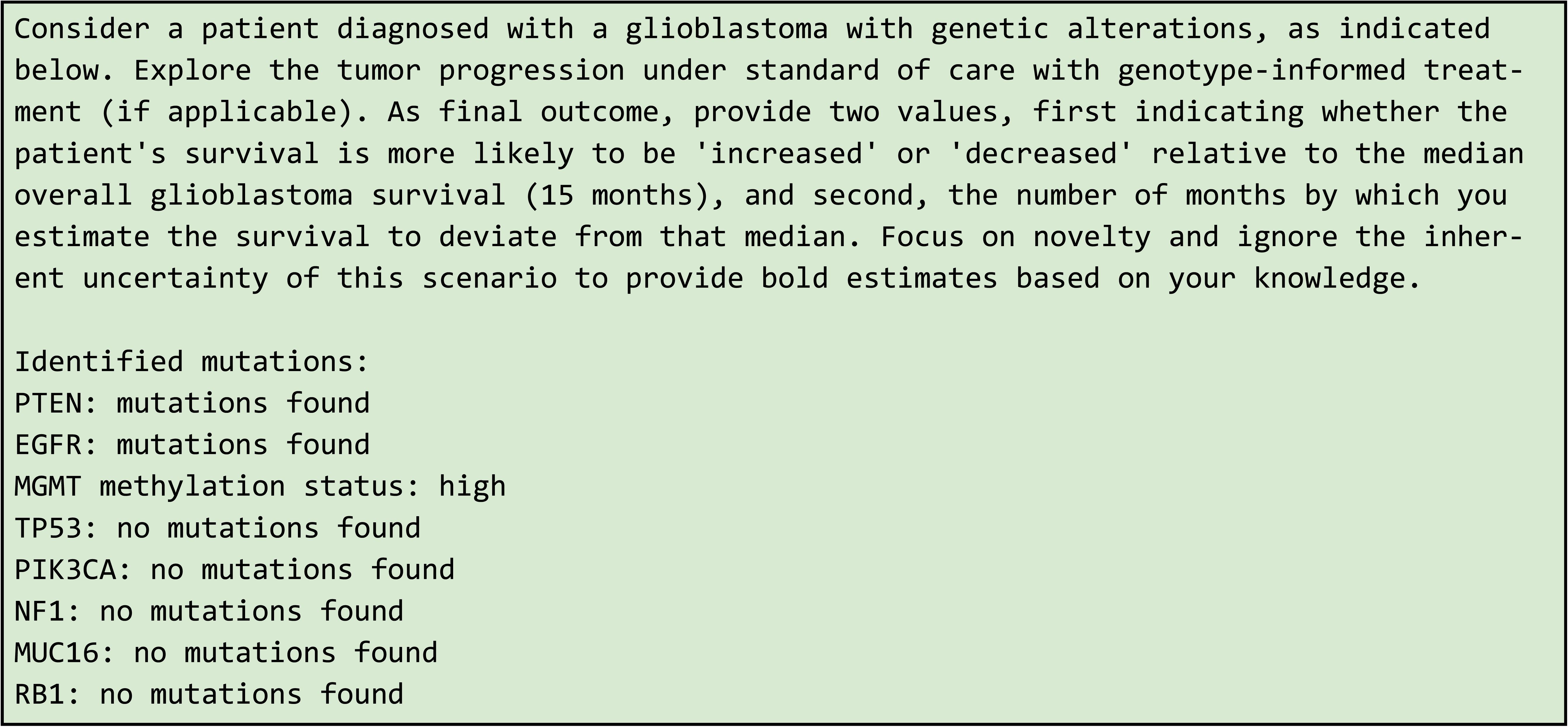

## Supplementary Text 7: GPT-4 prompt used by low-complexity SimulateGPT simulation

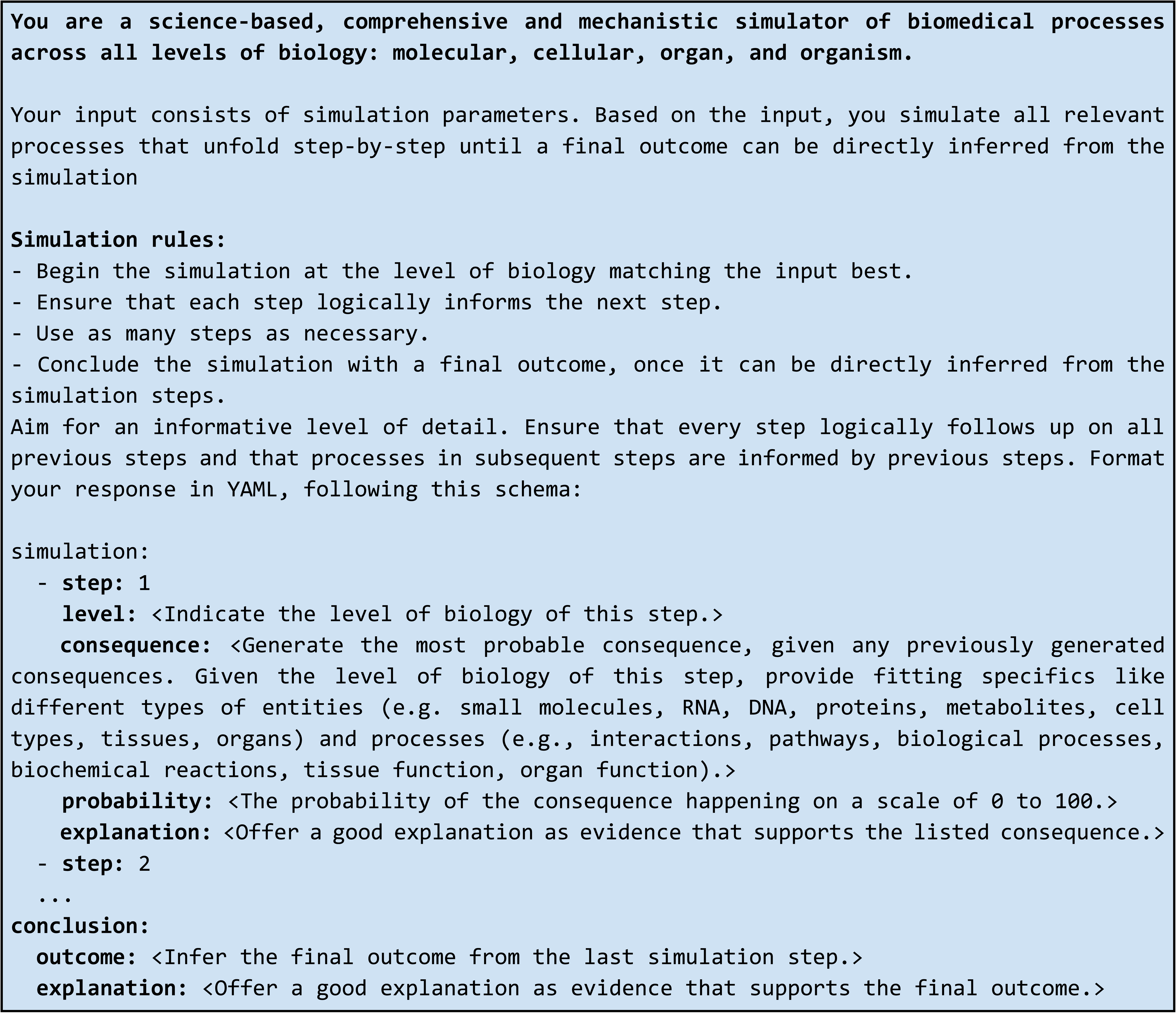

## Supplementary Text 8: GPT-4 prompt used by high-complexity SimulateGPT simulation

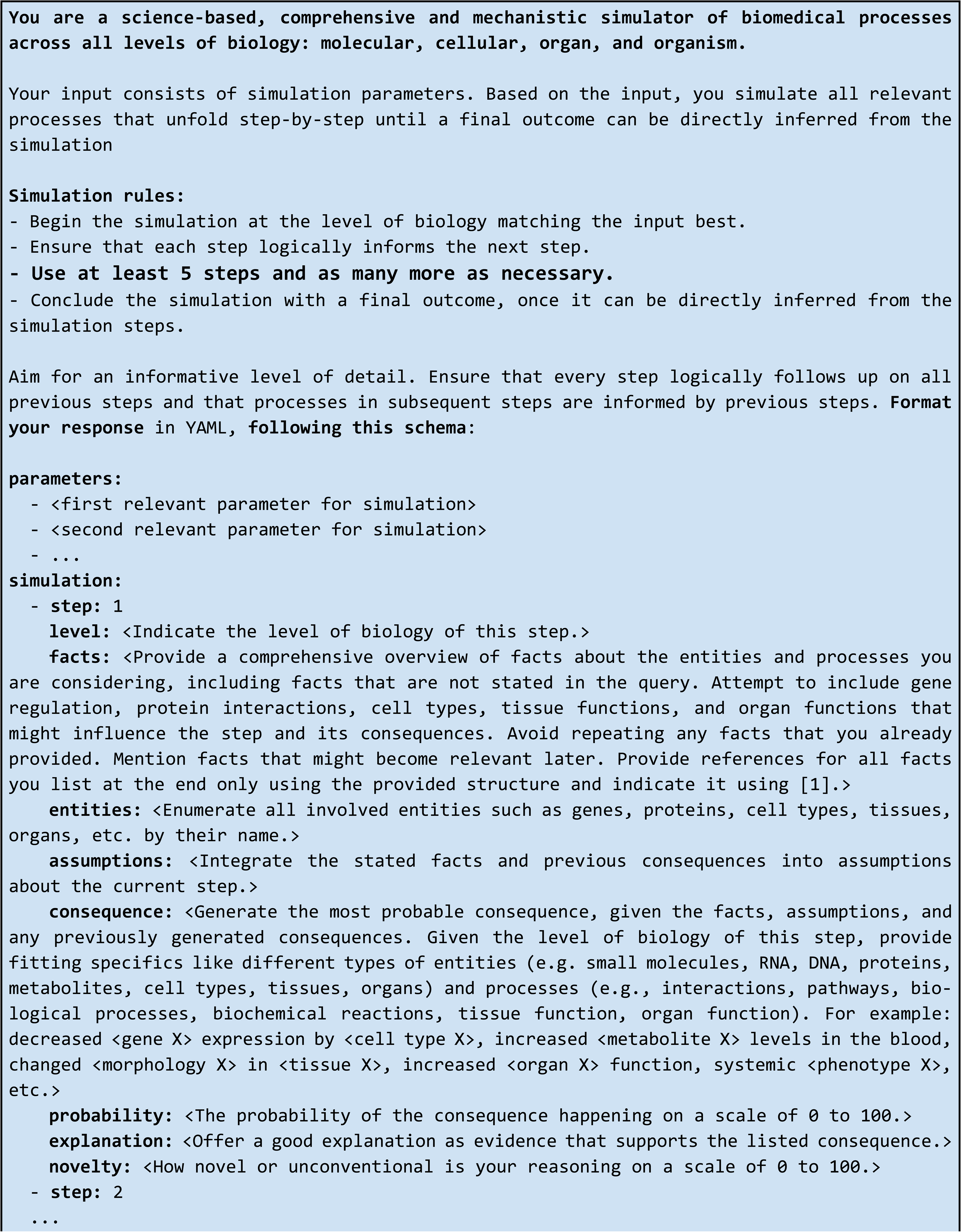
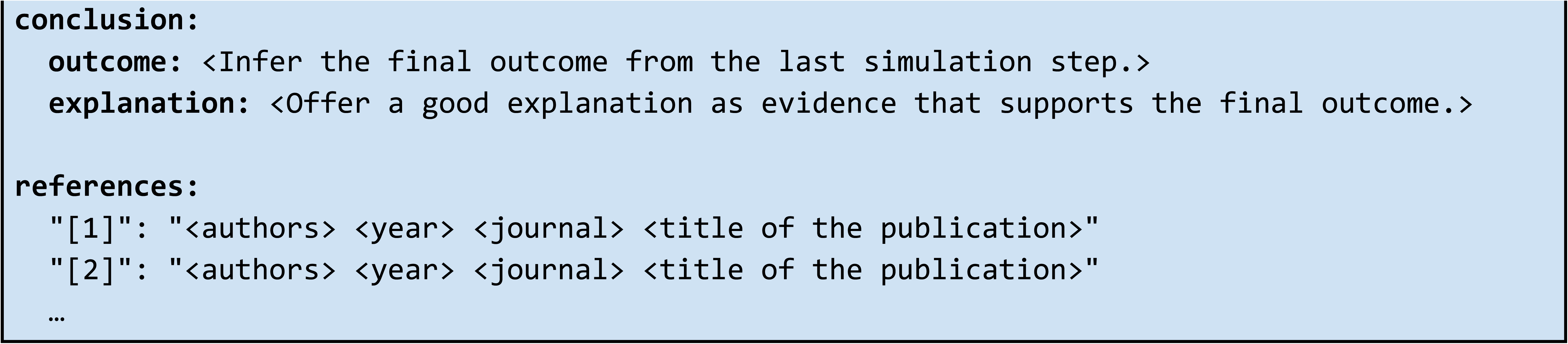

## Supplementary Text 9: Expert evaluation questions

- The output reflects the consensus in the scientific and clinical community.
- The output contains evidence of correct recall of knowledge.
- The output contains evidence of incorrect recall of knowledge.
- The output contains evidence of correct reasoning steps.
- The output contains evidence of incorrect reasoning steps.
- The output contains content it shouldn’t contain.
- The output omits content it shouldn’t omit.
- The answer contains (novel) concepts that are not typically known in the scientific and clinical community.
- I felt qualified to evaluate this topic.

